# An automatic domain-general error signal is shared across tasks and predicts confidence in different sensory modalities

**DOI:** 10.1101/2024.11.27.625775

**Authors:** Matthew Davidson, Sriraj Aiyer, Nick Yeung

## Abstract

Understanding the ability to self-evaluate decisions is an active area of research. This research has primarily focused on the neural correlates of self-evaluation during visual-tasks, and whether pre- or post-decisional neural correlates capture subjective confidence in that decision. This focus has been useful, yet also precludes an investigation of key every-day features of metacognitive self-evaluation: that decisions are rapid, must be evaluated without explicit feedback, and unfold in a multisensory world. These considerations lead us to hypothesise that an automatic domain-general metacognitive signal may be shared between sensory modalities, which we tested in the present study with multivariate decoding of electroencephalographic (EEG) data. Participants (N=21, 12 female) first performed a visual task with no request for self-evaluations of performance, prior to an auditory task that included rating decision confidence on each trial. A multivariate classifier trained to predict errors in the speeded visual-task generalised to predict errors in the subsequent non-speeded auditory discrimination. This generalisation was unique to classifiers trained on the visual response-locked data, and further predicted subjective confidence on the subsequent auditory task. This evidence of overlapping neural activity across the two tasks provides evidence for automatic encoding of confidence independent of any explicit request for metacognitive reports, and a shared basis for metacognitive evaluations across sensory modalities.

## Introduction

People can evaluate the quality of their decisions even in the absence of objective feedback, giving graded judgments of confidence (Henmon, 1911) and binary evaluations of accuracy (Rabbitt, 1968) that can correlate remarkably well with their objective performance. There is growing interest in the cognitive and neural mechanisms underpinning these “metacognitive” self-evaluations (Fleming & Frith, 2014), stimulated by evidence of the key role they play in adaptive behaviours such as information seeking (Desender et al., 2018; Pescetelli et al., 2021) and shared decision making (Bahrami et al., 2010; Silver et al., 2021).

The present research uses an EEG multivariate decoding approach to address three core questions regarding the mechanisms of metacognition. The first is the degree to which there is common coding of metacognitive evaluations across task domains. Whereas some evidence identifies dissociable correlates of confidence across different tasks, such as perceptual decisions versus memory retrieval (Baird et al., 2013; Fleming et al., 2014), other studies have found similar behavioural (Mazancieux et al., 2020) and neural markers (Rouault et al., 2023) associated with confidence in tasks as disparate as perceptual decisions and Sudoku puzzles (Su et al., 2022). Whether there is common encoding of confidence across domains is theoretically important, but also of practical significance given interest in exploiting neural signals of confidence to improve human decision making (Valeriani et al., 2017). Here we investigated the relatively understudied question of whether there are shared neural correlates across sensory modalities.

The second question we addressed is the relationship between graded judgments of confidence and binary judgments of response accuracy. There is an intuitive inverse relationship between judgments that a decision is correct versus judgments that one has made an error (Yeung & Summerfield, 2012). Consistent with this intuition, well-characterised markers of error detection that are observed in response-locked EEG data, the error-related negativity (ERN) and error positivity (Pe), also vary in amplitude in a graded manner with participants’ ratings of confidence in their decisions (Boldt & Yeung, 2015). However, several studies have documented EEG correlates of confidence in stimulus-locked waveforms (Gherman & Philiastides, 2015; Herding et al., 2019). Some studies have suggested further that post-response EEG activity may not index confidence despite the observed correlation (Rausch et al., 2020) and that the correlation itself may be a statistical artefact (Feuerriegel et al., 2022). These arguments align with the view that confidence is primarily determined by the strength of evidence accumulated to the point a decision is made (Kepecs & Mainen, 2012; Vickers & Packer, 1982), whereas post-decisional processing leading to error detection is largely restricted to tasks with speed pressure that induce “fast guess” errors (Rabbitt, 1968). Here we aimed to provide new evidence for a shared dependence of error detection and confidence judgments on post-response neural activity: We predicted that multivariate EEG activity associated with errors in a speeded (visual) task would generalise to predict graded variations in confidence in correct responses in an unspeeded (auditory) task.

Our final question was the degree to which metacognitive evaluations are an automatic and intrinsic part of decision making. Although some theories make this claim (Lee et al., 2023), evidence of “reactivity” (Double & Birney, 2019), such that asking participants to make metacognitive judgments alters their decision making (Petrusic & Baranski, 2003), suggests otherwise. The ubiquity of mind wandering (Killingsworth & Gilbert, 2010) likewise indicates imperfect and inconsistent self-monitoring (Jordano & Touron, 2018). In the present study, participants first performed the visual task with no request for (and, indeed, no mention of) self-evaluations of performance, prior to the auditory task in which participants were asked to report their decision confidence on each trial. Evidence of overlapping neural activity across the two tasks would provide evidence for automatic encoding of confidence independent of any explicit request for metacognitive reports.

## Method

### Participants

Twenty-one participants participated in this experiment (12 female, 20-35 years old, *M_age_* = 25), all with normal or corrected to normal vision. Three additional participants were recruited but excluded from subsequent analysis, for failing to adequately vary confidence ratings (single ratings when correct, *n* = 1) or poor EEG data quality (*n* = 2). All participants gave written informed consent and were paid for participation. The procedures were approved by the University of Oxford’s local ethics committee.

### Task and procedure

The experiment was designed for a within-participant comparison of neural activity following decisions on either a visual or auditory perceptual decision, with subjective confidence ratings only after the auditory task. We were motivated to explore whether post-decision error signals, in the absence of confidence judgements, could generalise to predict confidence formation in a separate modality. To this end, participants performed separate visual and auditory tasks in a fixed order while we recorded EEG data: Participants first completed blocks of a speeded visual discrimination task with no requested performance evaluations, before moving on to blocks of an auditory task for which they reported their subjective confidence after each decision (Figure 1).

**Figure 1.**
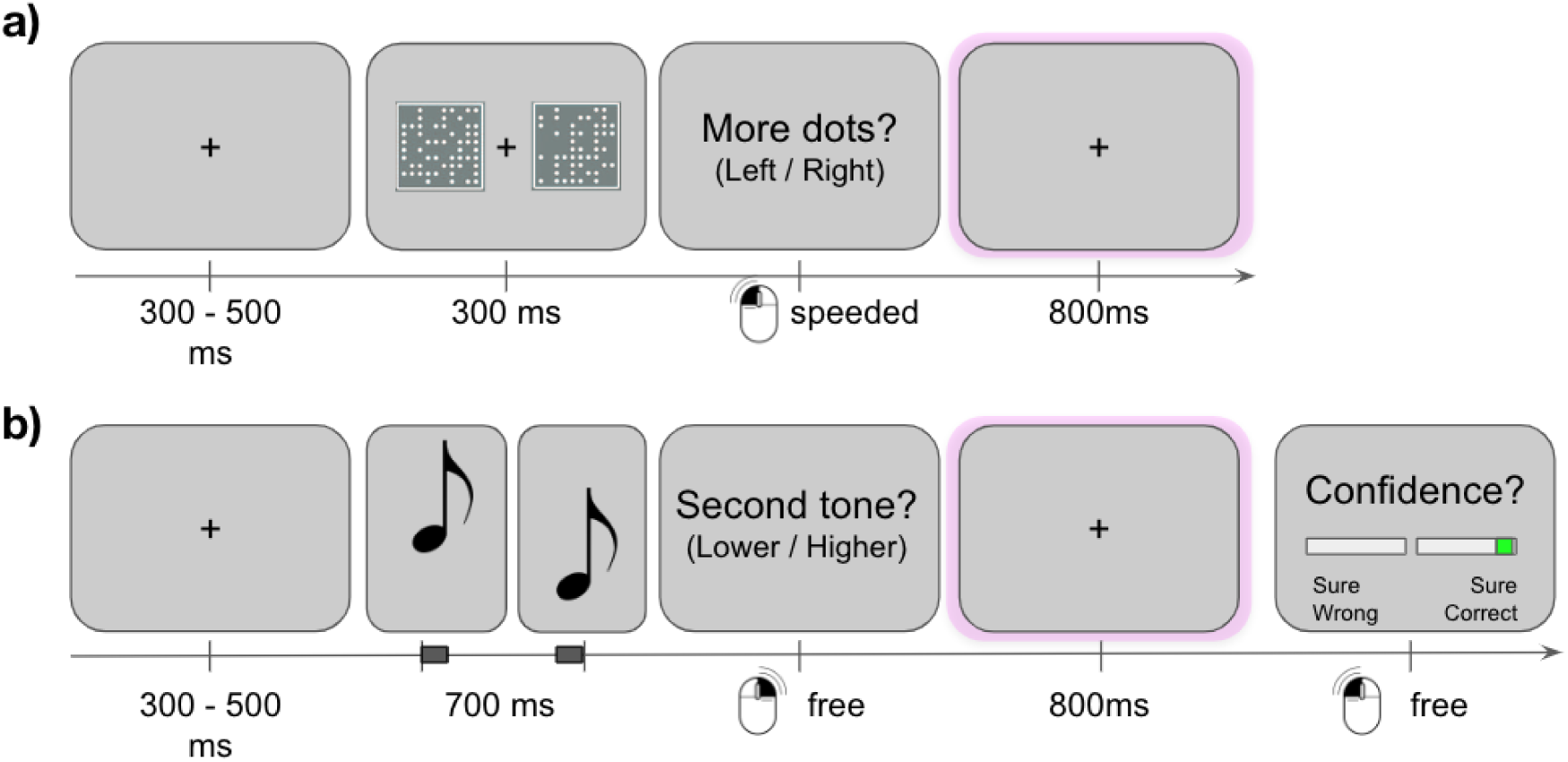
Experimental Paradigm. **a)** In the first half of each experiment, participants completed a speeded visual decision task. A fixed post-decision window was included to enable across-session comparison of neural activity. **b)** In the second half, participants completed an auditory decision task and additionally rated their confidence on each decision.

For the visual trials, participants were required to judge which of two square fields presented to the left or right of fixation contained more dots (300 ms duration, 400 dots total, left/right mouse click; Figure 1). Participants were not prompted to evaluate their trial wise performance at all during this stage—from their perspective, they simply performed an uninterrupted series of visual discriminations—and no written or verbal mention of confidence was provided to participants until after this phase was completed. The dot difference between fields was titrated per participant to approximate 84% accuracy, using a pre-experimental staircase procedure. During this staircase, 3 blocks of 30 trials were presented without feedback (one-up four-down adaptive staircase, increment 2 dots). After the staircase, the dot difference was fixed for the remaining 13 blocks of 30 trials of the visual experiment. If individual block accuracy exceeded 90%, or mean reaction times exceeded 2 seconds, participants were prompted to try to respond more quickly. Similarly, if block accuracy fell below 70% participants were instructed to respond more carefully. This emphasis on speed was used to enable the planned contrasts of neural activity on correct vs error trials, ensuring sufficient numbers of errors and also that many of these would be ‘fast guess’ errors that are typically associated with robust post-response error potentials (Scheffers & Coles, 2000). Importantly, after each response, we included a long intertrial interval (jittered with uniform distribution 1100 - 1300 ms), to facilitate the across-task decoding of response-locked neural signals related to the visual decision.

After completion of the visual portion of the experiment, participants were introduced to the auditory task and the additional confidence judgements that were required after each decision. On each trial in this phase, participants listened to two pure sine-tones in a quick sequence (100 msec tone, 500 msec gap, 100 msec tone) and were asked to judge whether the second tone of the sequence was lower or higher in pitch. The lower pitch was selected between 300 and 350 Hz on each trial. The frequency of the higher-pitch tones was set using an adaptive staircase procedure similar to the visual portion of the experiment, during which the ratio between low and high pitch tones was titrated to approximate 84% accuracy. Three blocks of 30 auditory calibration trials were presented during the staircase. After calibration, a further 10 blocks of 30 trials were completed at the fixed difficulty level. After each response on the auditory task, a fixed 800 msec interval was introduced before participants indicated their confidence in their previous decision. Confidence judgements were provided by clicking one of two slide bars, the first was labelled “Sure wrong” and ranged from 100% to 50% on the left of fixation, and second ranged from 50% to 100% “Sure correct” on the right of fixation (see Figure 1). There were no speed prompts between blocks in the auditory portion, confidence responses were not speeded, and after the response, the next trial began after a jittered 300-500 msec interval.

Stimuli were presented on a 17 inch monitor with a 60 Hz refresh rate using MATLAB (ver 2020) and Psychtoolbox routines (Brainard et al 1999). Dot fields were presented within a square frame (6°of visual angle wide) centred 5° from fixation (framewidth 0.1°). Each box was subdivided into a 20 x 20 grid, with square dots placed within grid locations selected from a random uniform distribution. Auditory stimuli were played at a comfortable volume over speakers placed either side of the monitor.

### EEG recording

Participants sat in an electrically shielded room and EEG was recorded at 1024 Hz from 64 active-electrodes corresponding to the 10-10 system (ActiCap: BrainProducts). An additional 6 electrodes were placed on the left/right canthi, above/below the right eye and on the left/right mastoid processes to monitor oculomotor activity and for offline re-referencing. All data was referenced online to FCz and re-referenced to the average of both mastoids during preprocessing. During preprocessing, EEG data were downsampled to 256 Hz, referenced to the average of both mastoids, and filtered between 0.1 and 30 Hz (zero-phase non-causal, 8449 point order, 0.1 Hz transition bandwidth). EEG were then epoched from - 500 ms to + 3500 ms relative to stimulus and response onsets. These datasets were then merged, and individual epochs inspected by eye for trial-rejection. On average, less than 4% of trials were identified for exclusion per participant. An Independent Components Analysis was performed to identify and remove blinks, oculomotor and other artefacts using the SASICA toolbox (Chaumon et al., 2015). After the ICA, automatic detection of channels with large kurtosis values (Z > 5) were spherically interpolated using nearest neighbours (average 2.7 channels per participant).

### Behavioural Data analysis

Accuracy and reaction times were calculated for each portion of the experimental task, excluding practice sessions. Auditory reaction times were calculated from the offset of the second tone stimulus. Confidence judgements were first z-scored within each participant to facilitate across-participant comparisons.

### EEG analysis

Event-related potentials were calculated aligned to visual stimulus onset, second auditory tone onset, and response onset per participant. Epochs were first linearly detrended, and then baseline corrected. Stimulus aligned epochs were baseline corrected relative to a 100 ms pre-stimulus window, and response-locked data were corrected using a −100 to −50 ms window in order to omit the interval containing the error-related negativity (ERN; cf. **Figure 3**). We note that the use of pre-response baseline may bias post-decision ERP amplitudes when systematic differences exist in the pre-response time-window (Feuerriegel and Bode 2022). In our dataset, we are not focused on interpreting the amplitude of the post-response waveform, but whether systematic whole-scalp variability encodes a metacognitive signal that is shared between tasks. This focus notwithstanding, we have also repeated our main analysis using a pre-stimulus baseline for comparison, and note our main results are not contingent on the pre-response baseline (e.g. crossmodal generalisation, Supplementary Figure 3). For our univariate ERP analysis, we calculated ERPs on correct and error trials, and when stratifying subjectively correct trials by subjective confidence in the auditory portion of the experiment. For this analysis, a median split of confidence values was used, owing to insufficient variation in confidence judgements to split by terciles of quartiles on a subset of participants (5/21).

**Figure 2.**
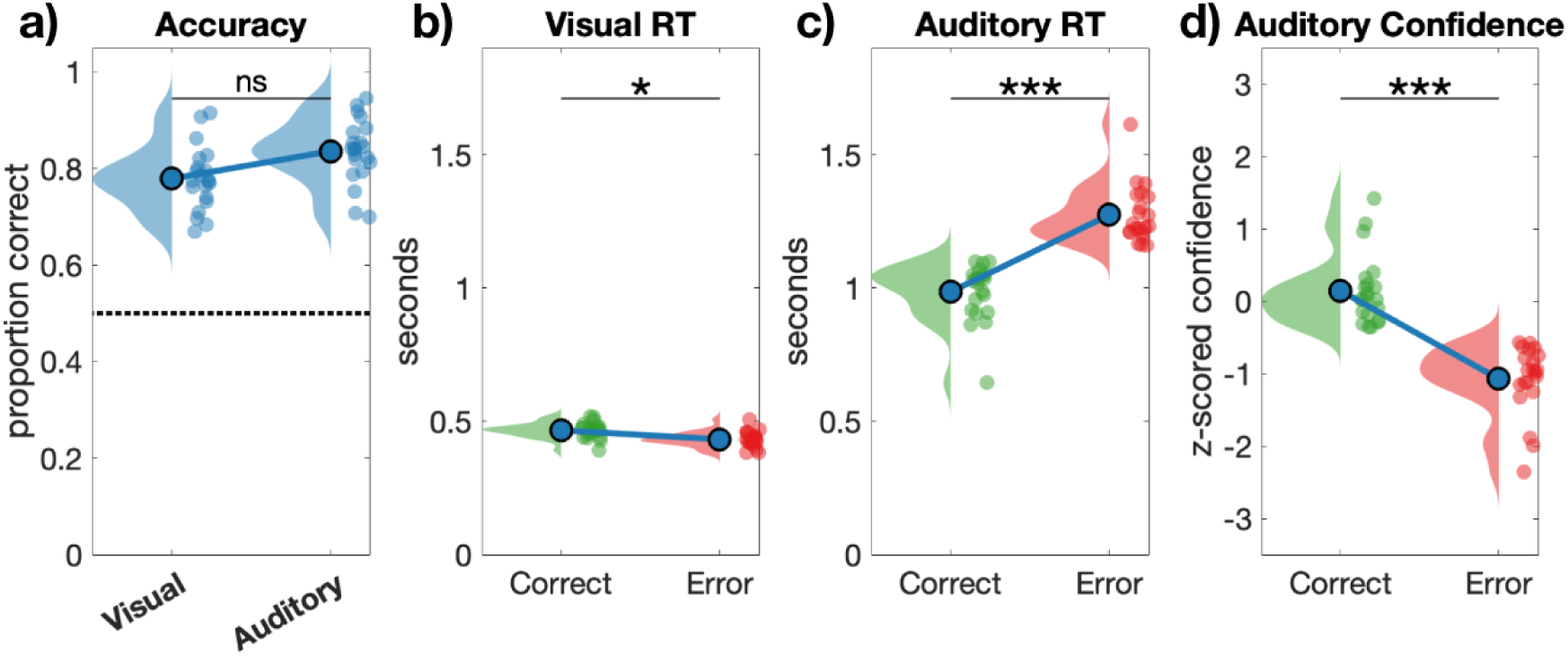
Behavioural results summary. **a)** Accuracy did not differ between the visual and auditory tasks. **b)** Reaction times were faster when committing errors on the speeded visual decision task, and **c)** reaction times were faster when correct on the auditory decision task. **d)** Confidence was calibrated with objective accuracy, with lower confidence overall on incorrect auditory decisions.

**Figure 3:**
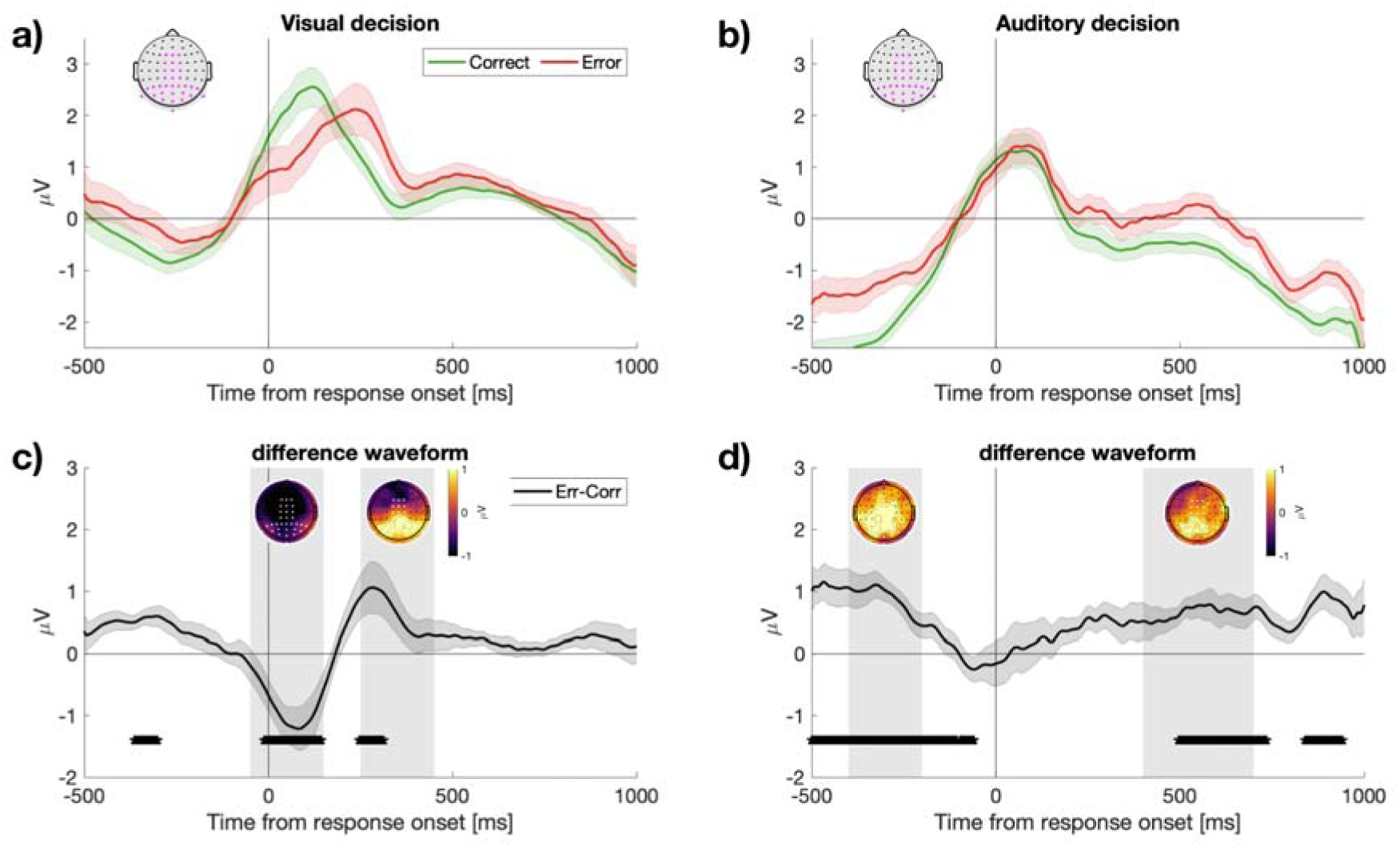
Response locked ERPs following a visual and auditory decision. **a)** ERPs time-locked to responses on the visual task, split by objective accuracy showing correct (green) and error (red) trials. **b)** ERPs time-locked to responses in the auditory task. In all panels, the ERPs are constructed from the electrodes marked in a-b in magenta. **c)** The difference waveform (error - correct) for visual responses. The inset plots show scalp topography for the average activity over the two temporal windows shaded in grey (−50 – 150 ms, and 250 – 450 ms). **d)** Difference waveform for auditory responses, with inset plots showing the scalp topography for the two temporal windows (−400 – −200 ms, and 400 – 700 ms). Asterisks denote *p* < .05 (uncorrected). **Supplementary Figure 1** displays the complementary stimulus-locked ERP data. **Supplementary Figure 2** displays the same response-locked data when using a pre-stimulus baseline.

For our main analysis, we trained a classifier to discriminate between correct and error trials using single-trial response-locked ERP waveforms (Parra et al., 2002; Boldt & Yeung, 2015). We were particularly motivated to test whether information in the Pe window (250 – 350 ms post response) would generalise to predict errors and confidence judgements in a separate auditory task. To robustly quantify Pe magnitude on single trials, we trained a multivariate classifier to distinguish between the response-locked ERPs generated on error and correct trials using the linear integration method (Parra et al., 2002). This method identifies the spatial topography that maximally discriminates between conditions of interest, which can be interpreted as a vector of weights that combine information from all electrode locations. Similar to conventional ERP analyses which improve signal-to-noise ratio by averaging over many channels, the linear integration method can improve the signal-to-noise ratio of single-trial data by combining the data across all electrodes via their weighted sum. This approach has previously been used to show that single-trial fluctuations in Pe amplitude predict subtle variations in subsequent confidence judgements (Boldt & Yeung, 2015). Here, we address whether a classifier trained in this manner can generalise to another modality, and whether single-trial Pe estimates are predictive of auditory confidence.

Our first analysis trained classifiers on the 250-350 ms window of post-response activity to align with the visual error related positivity (Pe; for similar see Steinhauser & Yeung, 2010; Boldt & Yeung, 2015). First, at the participant level, classifiers were trained using all error trials (Boldt & Yeung, 2015) and a matched size subset of correct trials which were selected at random from a uniform distribution (without replacement). As the selection of training trials could, in principle, have derived a different classifier, we repeated the selection of training trials over iterations of 5, 10, 20 and 50 validations. We note that participant level results were stable when using 5 or more iterations, and report results after calculating participant-level effects using 20 iterations. At the participant level, classifier performance was assessed by quantifying the proportion of trials that were correctly classified as errors or correct responses. **Figure 4** displays the group-level data after averaging across these participant-level effects. We display the probability of being labelled an error separately for both correct and error trials in the visual and auditory portion of the experiment, as well as the area under the ROC curve (AUC) when tested on all untrained trials.

**Figure 4.**
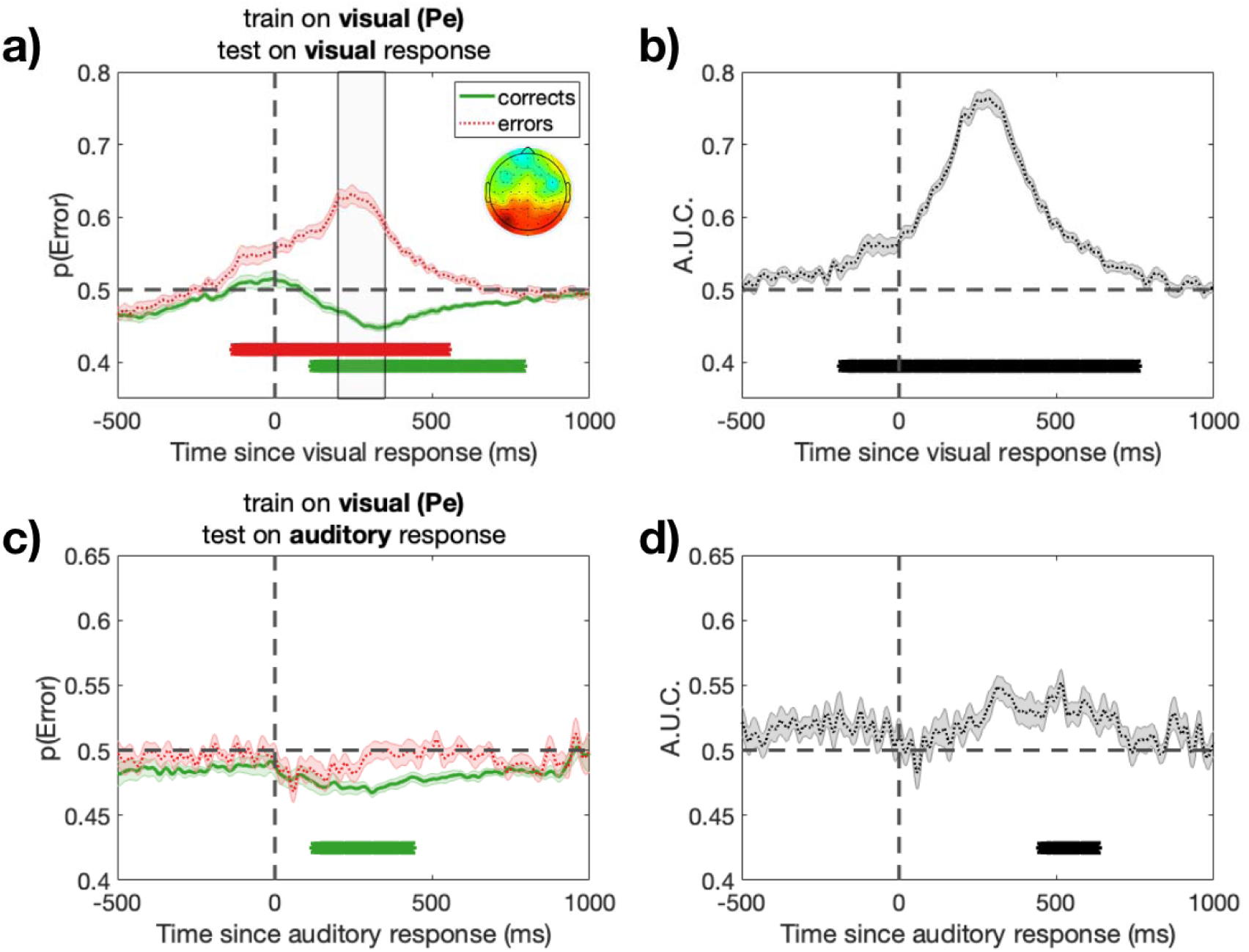
Temporal and crossmodal generalisation of a classifier trained to predict errors on a visual perceptual decision. **a)** Temporal generalisation of classifier performance when trained on the Pe window (250-350 ms, window shown in grey). The group-level weights assigned to each electrode are shown in the inset topoplot. Classifier performance significantly differs from chance on both error and correct trials (red and green asterisks respectively, *p*_cluster_ < .001) **b)** with high classification performance overall (black asterisks *p*_cluster_ <.001). **c)** The same classifier (trained on visual task data) generalises to predict correct trials on a later unspeeded auditory decision. **d)** Significant above chance classification occurs in the post-response window (*p*_cluster_ < .01).

We also examined the temporal-generalisation of classifier accuracy (King & Dehaene, 2014), and extended our training window and data-types to include all time points −500 ms – 1000 ms relative to both stimulus-onset and response-onset locked ERP waveforms. This analysis was performed to visualise whether error and correct trials in an auditory task could also be predicted from any stimulus-locked activity, and whether activity in the response-locked Pe window was unique in this regard. This analysis used a sliding window approach creating classifiers based on 50 msec windows of activity progressing in steps of 25 msec. Classifier training and test data were selected as above. The temporal-generalisation of each classifier was assessed by applying the topography obtained from each 50 msec window to all time-points, before averaging AUC values within each 50 ms training window for analysis and visualisation (**Figure 5**). As classifiers were trained and tested on all time-points this creates a 2D performance matrix.

**Figure 5.**
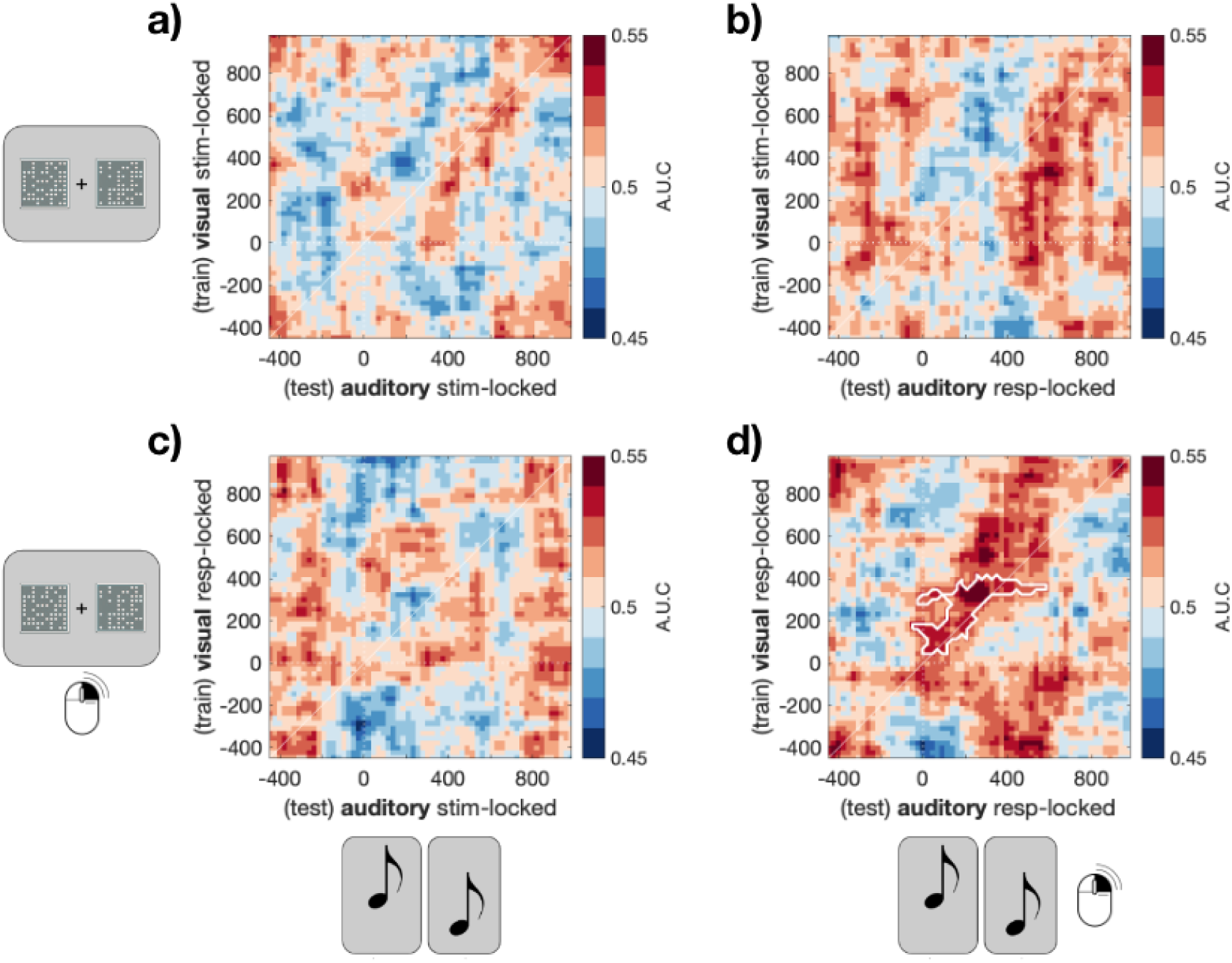
Crossmodal and temporal generalisation of error classification. Each panel shows the temporal generalisation matrix of multivariate classifiers trained on either the visual stimulus **(a,b)**- or visual response locked ERP **(c,d)**, and tested on either the auditory stimulus-locked **(a,c)** or response-locked ERP (**b,d)**. Only a classifier trained on the visual response-locked ERP data generalises to predict errors in a subsequent auditory task **(d)** (white lines denote boundaries, *p*_cluster_ < .001 corrected for multiple comparisons (Maris & Oostenveld, 2007)).

Contiguous significant clusters of activity were first identified at an alpha level of *p* < .05 (two-tailed, uncorrected, *t*-test against chance), and the absolute sum of *t* values was retained per cluster for comparison with a null-distribution of cluster-level test statistics. To create the null-distribution the temporal-generalisation of classifier performance was repeated on trials with randomised condition labels (1000 permutations). The maximum cluster-level test-statistic was retained per permutation, as described above, and the originally observed cluster-level statistics were considered significant when exceeding the bottom 95th percentile of this null distribution (e.g. *p*_cluster_ < .05) The results of this analysis are displayed in **Figure 5**.

Finally, after demonstrating that the Pe window classifier could generalise to predict errors in a crossmodal task, we investigated whether classifier output would correlate with single-trial confidence ratings. For this analysis, we restricted our focus to only include subjectively correct trials – trials in which participants rated their confidence between 50 – 100% “Sure Correct”. This subset excludes trials in which participants may have changed their mind after their initial response and detected errors which may otherwise confound overall classifier performance. Next, we applied the classifier weights derived from the visual Pe window to all auditory response-locked data using a sliding window approach (50 ms window, 5 ms step size), retaining a score per time-point across all trials for evidence of an objectively correct or error response (Bernoulli probability distribution, ranging from 0 to 1 respectively). All trials were then correlated with their respective confidence ratings, and the correlation visualised per 5 ms step in our sliding window approach. This analysis effectively quantifies whether (and when) confidence judgements correlate with neural activity that resembles visual error detection during the auditory portion of the task. The results of these analyses are shown in **Figure 6**. As above, we corrected for multiple comparisons in the correlation time-series using a non-parametric cluster-based correction (Maris & Oostenveld, 2007). Temporally contiguous clusters were first detected (at *p* < .05, two-tailed, uncorrected) before the summed cluster-level test-statistic was retained for comparison to the null distribution. The null distribution was created by shuffling condition labels 2000 times, before retaining the maximum sum of cluster-level test statistics of each permutation. We regarded the observed cluster-level test-statistic to be significant if it exceeded the bottom 95% of the null distribution of the summed statistics (e.g. *p*_cluster_ < .05).

**Figure 6.**
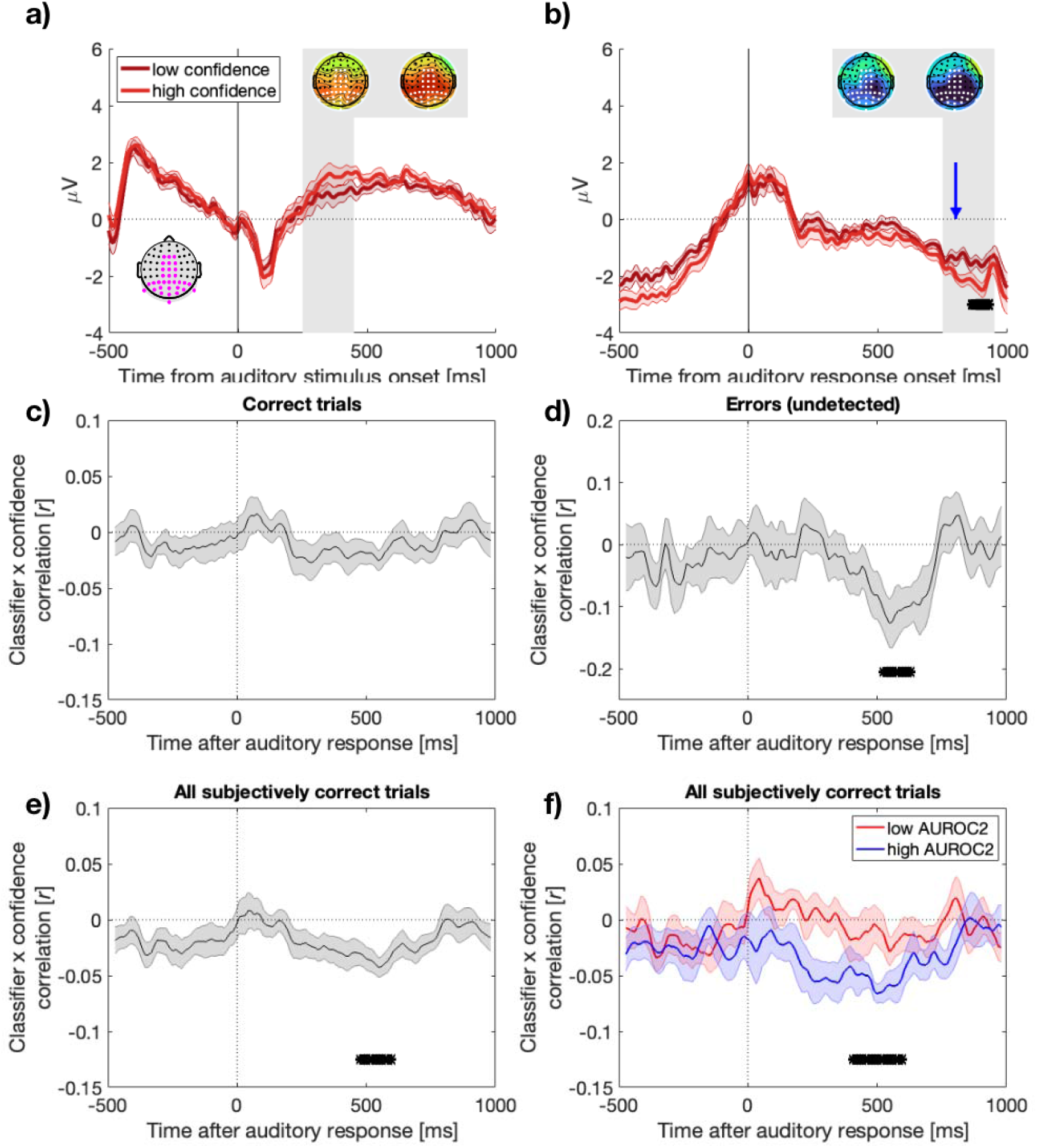
Visual error detection underpins auditory confidence. **a-b)** Low and high-confidence ERP waveforms defined by median split per participant, within the restricted subset of subjectively correct trials only. a) Displays auditory-stimulus locked waveforms, b) displays auditory response-locked waveforms. Channels used to compute ERPs are shown in the inset (grey, magenta). Topoplots of average activity within grey shaded windows of interest (250 – 450 ms) and 750 – 950 ms post-stimulus and response, respectively) are included. Left topoplots show low-confidence trials, Right topoplots show high-confidence trials. The vertical blue line indicates the onset of the confidence response screen. **c)** Single-trial correlation between error classification and confidence judgements (see Methods). The topographic weights from a classifier trained exclusively on the Pe window following responses to a visual stimulus are shown in the inset (arbitrary units). These weights were applied to the activity in all auditory response-locked waveforms using a sliding window approach, restricting to subjectively correct trials only. The classifier linearly integrates activity over all electrodes to produce a probability for the likelihood of a visual error. **c)** Shows the classifier did not correlate with confidence on objectively correct trials. The negative correlation in **d)** indicates that confidence judgements on the auditory task decrease as auditory-response locked neural activity resembles visual errors, even when participants were unaware of these errors and rated confidence in the range of subjectively correct. **e)** Displays the result from all subjectively correct trials. **f)** The same result as **e)** after median-split of the participants based on type-2 AUC (metacognitive sensitivity). Asterisks in all plots denote *p*_cluster_ <.05.

## Results

### Behavioural data

Participants responded to a speeded visual perception task indicating which of two locations contained a larger number of dots, as well as an auditory decision task which additionally prompted unspeeded confidence judgements. **Figure 1** displays a schematic of the trial structure, and **Figure 2** shows the behavioural results.

Both the visual and auditory portions of the experiment began with an adaptive staircase procedure to calibrate performance to approximately 84% accuracy. As a result, there was no significant difference in accuracy on the two tasks (*p* = .06). However, the relative speed of correct vs. error responses differed across tasks. Thus, as is typically observed in speeded tasks with many fast guess errors, visual task reaction times were shorter on error trials (*M* = 0.43 s, *SD* = 0.12), than they were on correct trials (*M* = 0.46 s, *SD* = .08; *t*(20)= 2.70, *p* = .014, *d* = 0.59). In contrast, when performing the unspeeded auditory portion of the experiment, reaction times were longer on error trials (Errors *M* = 1.27 s, *SD* = 0.37; Corrects *M* = 0.99 s, *SD* = 0.29; *t*(20)= −6.18, *p* < .001, *d* = 1.35). Confidence judgements were also lower on error trials (Corrects *M* = 0.15, *SD* = .07; Errors *M* = −1.07, *SD* = .92; *t*(20) = 5.81, *p* < .001, *d =* 1.27). Reflecting that incorrect responses in the unspeeded auditory task tended to be “data limited errors” that reflected the difficulty of the perceptual discrimination rather than the occurrence of fast guesses (Scheffers & Coles, 2000), participants’ confidence ratings fell in the range ‘guess’ to ‘sure correct’ on 76.67% of trials (SD = 22.78, range 9.1% – 97.5%), indicating a predominant failure of participants to detect those errors.

### Univariate ERP data

We proceeded by investigating the post-decision event-related potentials. **Figure 3** displays a summary of the response-locked univariate ERP data. Consistent with prior research, when computing the difference waveform between error and correct trials in the visual task, we observed a pronounced fronto-central negativity beginning around the time of the response and peaking shortly after, followed by a positive component approximately 200-400 ms post-response that was maximal over posterior scalp sites. These components align with the error-related negativity (ERN), and error positivity (Pe), implicated in previous investigations of post-decision confidence formation. The amplitude of the Pe in particular has been shown to correlate with subjectively reported confidence (Boldt & Yeung, 2015), and whether neural-activity in this window generalises to predict auditory decision making is our main focus. For comparison, the right-hand panels of **Figure 3** plot the corresponding waveforms for the auditory task. There is no prominent ERN component, but a Pe is apparent as an increased positive voltage for the error-trial waveform over posterior sites. Compared with the visual-task Pe, however, the auditory-task Pe has a markedly slower onset and longer duration.

### Single-trial multivariate decoding

Our univariate analysis revealed pronounced differences in ERP activity when comparing correct and error trials in both the visual and auditory tasks. Notably, the magnitude and morphology of these responses differed based on the modality of the task (see **Figure 3 c-d**). However, of key interest was whether we would nevertheless find evidence of shared neural signatures of metacognitive monitoring in the two tasks. To this end, our next analysis investigated whether a multivariate classifier trained to predict error responses in the visual task would generalise to correctly predict errors in an auditory decision.

For this analysis, we focused on the Pe in a window from 250-350 msec post-response, in line with our previous work (Boldt & Yeung, 2015; Steinhauser & Yeung, 2010). We trained our classifier to distinguish error and correct trials of the visual task to derive a spatial weighting of electrodes that, when convolved with observed scalp voltages, maximally distinguishes those trial subsets. The resulting weighting can then be applied to other data outside the training set, with the output being a value for each timepoint that corresponds to the estimated probability that the trial belongs to the target category (here: errors). The temporal generalisation of the classifier as applied to the response-locked EEG data from the visual task is shown in **Figure 4a**, for correct and error trials separately. The classifier shows significant classification of both errors and correct responses for sustained periods in the post-response period, assigning the former a p(Error) value greater than 0.5 and the latter a value less than 0.5. When tested on all trials, the area under the ROC curve is displayed in **4b.** Peak classification accuracy occurs within the training-window, as expected, but above chance decoding accuracy is also clear for earlier and later periods around the time of response, from approximately −435 to 765 msec.

Importantly, we next tested the visual classifier on the auditory response-locked data, wherein none of the all auditory trials were used in training, and observed significant above chance classification over the period 265 to msec after the response (see **Figure 4c,d**). These findings suggest overlapping post-decisional neural markers of metacognition across the visual and auditory tasks. The correspondence between the timing of significant cross-classification (**Figure 4d**) and of the auditory task Pe (**Figure 3b,d**) is consistent with this component being the shared neural correlate of metacognitive evaluation. The time-limited nature of cross-classification, which dissipates towards the end of the response-locked epoch, rules out the results being caused by stimulus-locked activity contaminating the response-locked baseline period we use (cf. Feuerriegel et al. 2022), any effect of which should persist throughout the epoch. Confirming this, similar results are observed when we reanalyse the data using a pre-stimulus rather than pre-response baseline—see **Supplementary Figure 3**. Notably, classification primarily reflected assignment of P(Error) values lower than 0.5 to correct auditory task trials rather than values above 0.5 to errors (**Figure 4c**). For errors, the P(Error) classifier value did not significantly exceed 0.5 at any time point. These findings are a first indication that the classifier is generalising to identify variations in confidence in correct responses, rather than only participants detecting their errors, a theme explored further below.

### Temporal generalisation of crossmodal classifier accuracy

**Figure 4** demonstrates that a classifier trained on the Pe window following a visual decision can generalise to classify response accuracy in a later, unspeeded auditory decision. The time-course of changing classifier accuracy extending beyond the training window (cf. Figure 4b**,d**) also demonstrates the temporal generalisation of this crossmodal classification. Our next analysis formally quantified the extent of this temporal generalisation by comparing all stimulus locked and response locked time-points, using the temporal-generalisation method (King & Dehaene, 2014).

For this analysis, we repeated the same procedure as described above but now using a variety of time-windows in both stimulus- and response-locked epochs for both training and testing. With this approach, we tested whether the generalised prediction of crossmodal error was unique to the response-locked Pe window, or would also be observed in earlier stimulus-locked differences in ERP amplitude, and further whether cross-classification would be possible between stimulus- and response-locked waveforms. Figure 5 shows the results of this analysis. Classifiers trained on stimulus-locked ERP waveforms from the visual task did not lead to above-chance decoding accuracy for auditory decisions, either when tested on stimulus-locked (Figure 5a) or response-locked waveforms (Figure 5b). Nor did classifiers trained on visual task response-locked waveforms generalise to classify auditory task trials based on stimulus-locked EEG activity (Figure 5c). Significant cross-classification was only apparent in the response-locked waveforms (Figure 5d). This result extends the basic classification analysis reported above (Figure 4) to show that significant cross-task classification is evident for a range of training and testing timepoints across the window from 0 to 600 msec post-response. Cross-classification is primarily observed along the diagonal. The limited temporal generalisation (i.e., classifiers trained on one timepoint do not robustly cross-classify data from distant timepoints) can be indicative of a dynamic sequence of neural representations (King & Dehaene, 2014).

### Classifier scores correlate with single-trial confidence ratings

Our final analysis provided the critical evaluation of whether our classifier trained to distinguish correct from error responses in a visual task would generalise further to predict subtle fluctuations in confidence judgements in the auditory task. We hypothesised that as well as showing above chance decoding accuracy (cf. Figure 4d), classifier scores would also negatively correlate with confidence judgements in a time-dependent manner (negatively because the classifier output corresponds to P(Error), which should vary inversely with participants’ estimates that their responses are correct). Evidence consistent with this prediction would demonstrate that a supramodal error signal (formed in the absence of explicit confidence reports) also underpins the strength of later confidence judgements in a crossmodal task. For this analysis, we applied the classifier trained on the visual Pe window to all time-points within the auditory response-locked data, and calculated the correlation using a sliding window approach (see Methods). We were specifically interested in variations in participants’ confidence in correct decisions, and therefore restricted analysis to trials in which participants rated their confidence as above 50% (i.e., excluding trials on which participants detected errors). **Figure 6** displays a summary of the results.

**Figure 6** plots ERP waveforms when subjectively-correct trials are median split into low vs. high confidence subsets and the data are aligned to stimulus or response-onset. The waveforms show only modest differences, with small and non-significant differences in the time range of the Pe (where we might expect lower confidence trials to be associated with a Pe-like positivity). Small differences in ERP amplitude within subjectively correct trials emerged close to the time of providing a subjective confidence judgement. Nevertheless, as predicted, when applied to these subjectively-correct trials, the scalp topography used to distinguish error trials in the visual portion of the experiment negatively correlated with confidence judgements in the auditory portion of the task (**Figure 6d,e**). Thus, the EEG-derived estimate of P(Error) for the time window from around 500-600 msec after the response predicted participants subsequently-reported confidence. As with cross-classification of auditory task accuracy (**Figure 4d**), the time-course of this correlation tracked the morphology of the Pe component. Ruling out that the correlation reflected contamination of our response-locked baseline by stimulus-locked activity, the correlation was clearly time limited rather than sustained to the end of the analysed epoch.

As further evidence that the classifier was sensitive to a shared neural marker of error detection and subjective confidence, we repeated the analysis in **Figure 6e** after performing a median-split of our participant sample based on metacognitive sensitivity. We quantified metacognitive-sensitivity (type 2 performance) as the area under the receiver operating characteristic curve per participant, which captures the fidelity of confidence judgements to predict objective performance. **Figure 6f** displays that the group-level result of a negative correlation between classifier performance and confidence values is primarily driven by the subset of participants with relatively higher metacognitive sensitivity, whose confidence values were superior indicators of objective accuracy.

## Discussion

Our EEG decoding and generalisation analyses provide evidence for shared neural correlates of implicit error detection between visual and auditory tasks. We trained a multivariate classifier to distinguish the EEG activity observed following errors versus correct responses in a visual perceptual task. We found, as predicted, that this classifier generalised to distinguish correct responses from errors in a separate auditory pitch-comparison task, and further to predict variations in participants’ reported decision confidence in that task.

A first implication of our results is that there are at least partially overlapping markers of metacognitive evaluations across tasks involving different sensory modalities. This finding adds to a growing corpus of neuroimaging evidence for common coding of confidence across different kinds of decisions (Fernandez-Vargas et al., 2021; Rouault et al., 2023; Su et al., 2022), a finding that has also been reported in other species (Masset et al., 2020). This evidence notwithstanding, several studies have also reported dissociations, for example between neural mechanisms supporting metacognitive evaluations in perception and memory (Baird et al., 2013; Fleming et al., 2014). Indeed, evidence of both overlap and dissociations have sometimes been reported in the same study (Morales et al., 2018; Qiu et al., 2018; Su et al., 2022), leading to the idea of hierarchical structure whereby domain-general confidence signals in frontoparietal circuits are transformed into content-rich, task-dependent representations in anterior prefrontal cortex that subserve adaptive, context-specific control of behaviour. Within this framework, the purpose of domain-general confidence representations might lie in allowing generalisation of adaptive behaviours (such as information seeking when unsure) across different cognitive tasks (Desender et al., 2018; Kornell et al., 2007), as well as in allowing adjudication between different kinds of decisions based on their relative reliability (de Gardelle & Mamassian, 2014). Meanwhile, there is practical value in identifying domain-general markers of confidence given their potential use in improving human decision making via brain-computer interfaces (Valeriani et al., 2017). Our contribution is to demonstrate this domain-generality using EEG decoding across tasks in different sensory modalities.

Our second, related contribution is to locate a shared basis of metacognitive evaluations in post-decisional processes: Our multivariate decoding analysis focused on the time-window of the response-locked Pe component, which has previously been shown to vary with graded evaluations of error likelihood (Steinhauser & Yeung, 2010) and confidence (Boldt & Yeung, 2015) in the visual task used here. Error detection inherently depends on continued processing after an initial decision that leads to a change of mind (Rabbitt, 1968). The sensitivity of confidence judgments to post-decisional processing is more controversial: Although proposed by some theories (Pleskac & Busemeyer, 2010) and supported by various lines of evidence (Charles & Yeung, 2019; van den Berg et al., 2016; Yu et al., 2015), counter-arguments and counter-evidence exist (Feuerriegel et al., 2022; Rausch et al., 2020). Here we show that the association between post-decisional EEG activity and confidence is not restricted to speeded tasks with frequent “fast guess” errors: our auditory task was unspeeded and its errors were typically slow responses. Nor can the correlation we observe between Pe amplitude and confidence be explained in terms of artifacts arising from stimulus-locked activity contaminating a response-locked baseline (Feuerriegel et al., 2022) because, as also shown previously (Boldt & Yeung, 2015), the relationship is time-limited and holds even in analyses using a pre-stimulus baseline. Indeed, we found reliable cross-task decoding only in response-locked data, with no significant generalised decoding in analyses of stimulus-locked data. The lack of stimulus-locked decoding of confidence is initially surprising given longstanding evidence linking confidence to the amplitude of the stimulus-locked P3 component (Gherman & Philiastides, 2015; Herding et al., 2019; Hillyard et al., 1971). However, our approach may not be optimal for exploring the confidence-P3 relationship, for which an approach based on decoding confidence directly (rather than indirectly via initially decoding the contrast between errors and correct trials) might produce more positive results. That is a potential direction for future research. Our contribution here is to demonstrate overlap between neural correlates of error monitoring and confidence judgments via a shared dependence, across tasks, on post-decisional processing.

The third question addressed in our study is the degree to which metacognitive evaluations are an automatic and inherent part of the decision making process. Although often assumed, and sometimes explicitly theorised (Lee et al., 2023), there is little direct study of how often thoughts are metacognitive (Jordano & Touron, 2018). Error-related EEG activity was originally reported in tasks that did not ask participants to make any explicit performance evaluation (Falkenstein, 1990) and it shows only small modulation when explicit reports are requested (Grützmann et al., 2014), suggesting that error detection proceeds automatically and implicitly in typical cognitive tasks. Here we have shown that error-related signals observed in a (visual) task with no explicit performance monitoring component generalise to predict participants’ confidence reports in a separate (auditory) task. To the degree that these findings indicate that the Pe component is best characterised as a graded signal that reflects subtle variation in confidence, not simply a binary classification of decisions as correct versus incorrect, they suggest that people generate graded evaluations of confidence even when there is no requirement for them to do so.

Future research may wish to leverage this evidence of a shared domain-general metacognitive signal, yet critical questions remain. While we have focused on two nominally distinct tasks (visual speeded vs auditory unspeeded), both required a binary two alternative forced-choice response. Extensions beyond binary tasks, which also vary response protocols could provide further evidence of an automatic and generalisable metacognitive self-evaluation signal.

## Supplementary Figures

**Supplementary Figure 1.**
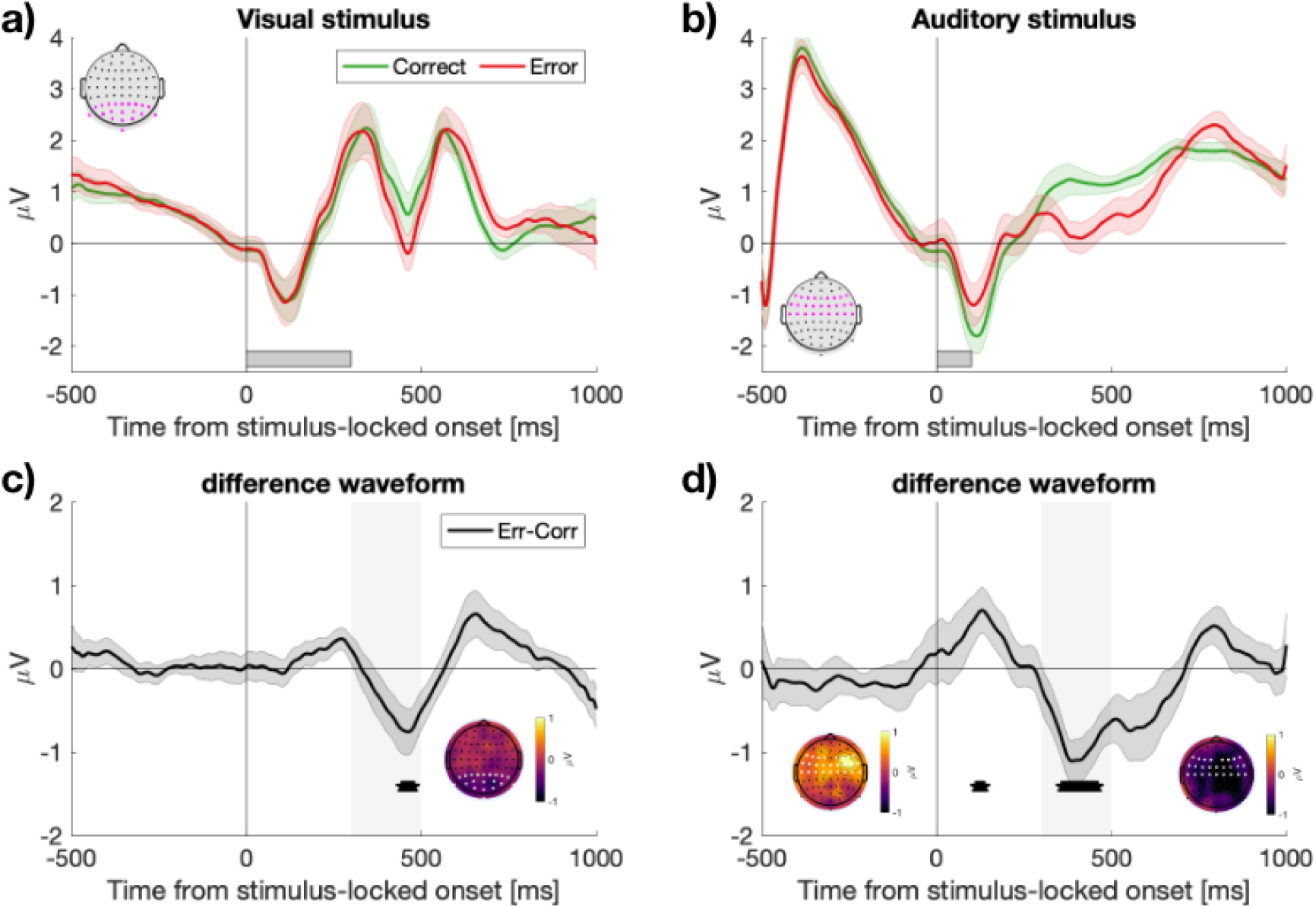
Stimulus-locked ERPs following a visual and auditory stimulus. **a)** ERPs time-locked to the onset of the dot display in the visual task, split by objective accuracy showing correct (green) and error (red) trials. The dots were on screen for 300 ms, as indicated by the horizontal grey bar. **b)** ERPs time-locked to the onset of the second auditory tone in the auditory task. In all panels, the ERPs are constructed from the electrodes marked in a-b in magenta. **c)** The difference waveform (error -correct) for visual ERPs in a). **d)** Difference waveform for auditory ERPs in b). The inset topoplots show average activity over the two temporal windows shaded in grey (300 – 500 ms). Black asterisks denote *p* < .05 (uncorrected).

**Supplementary Figure 2.**
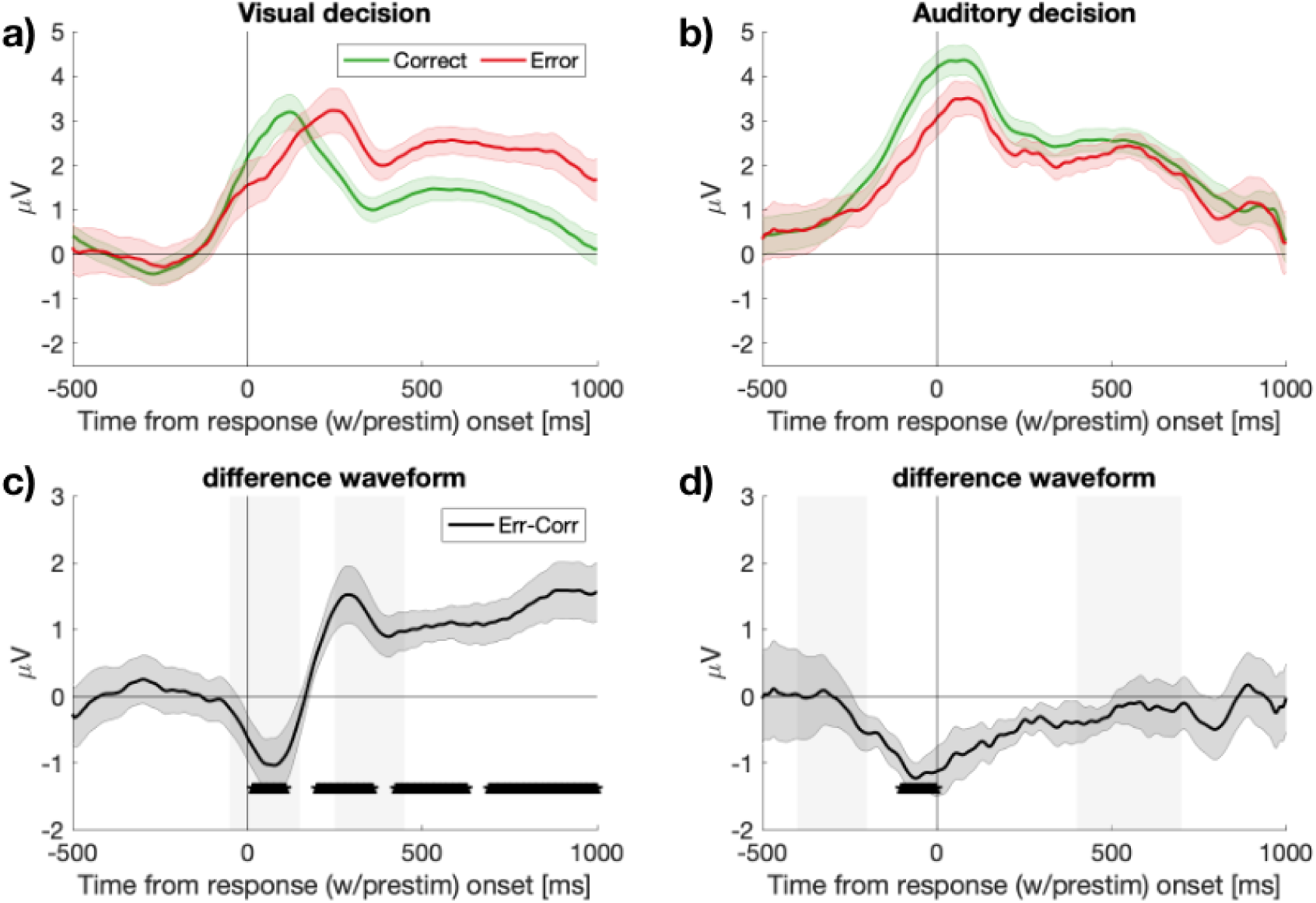
Response-locked ERPs following a visual and auditory stimulus, using pre-stimulus baseline.

**Supplementary Figure 3.**
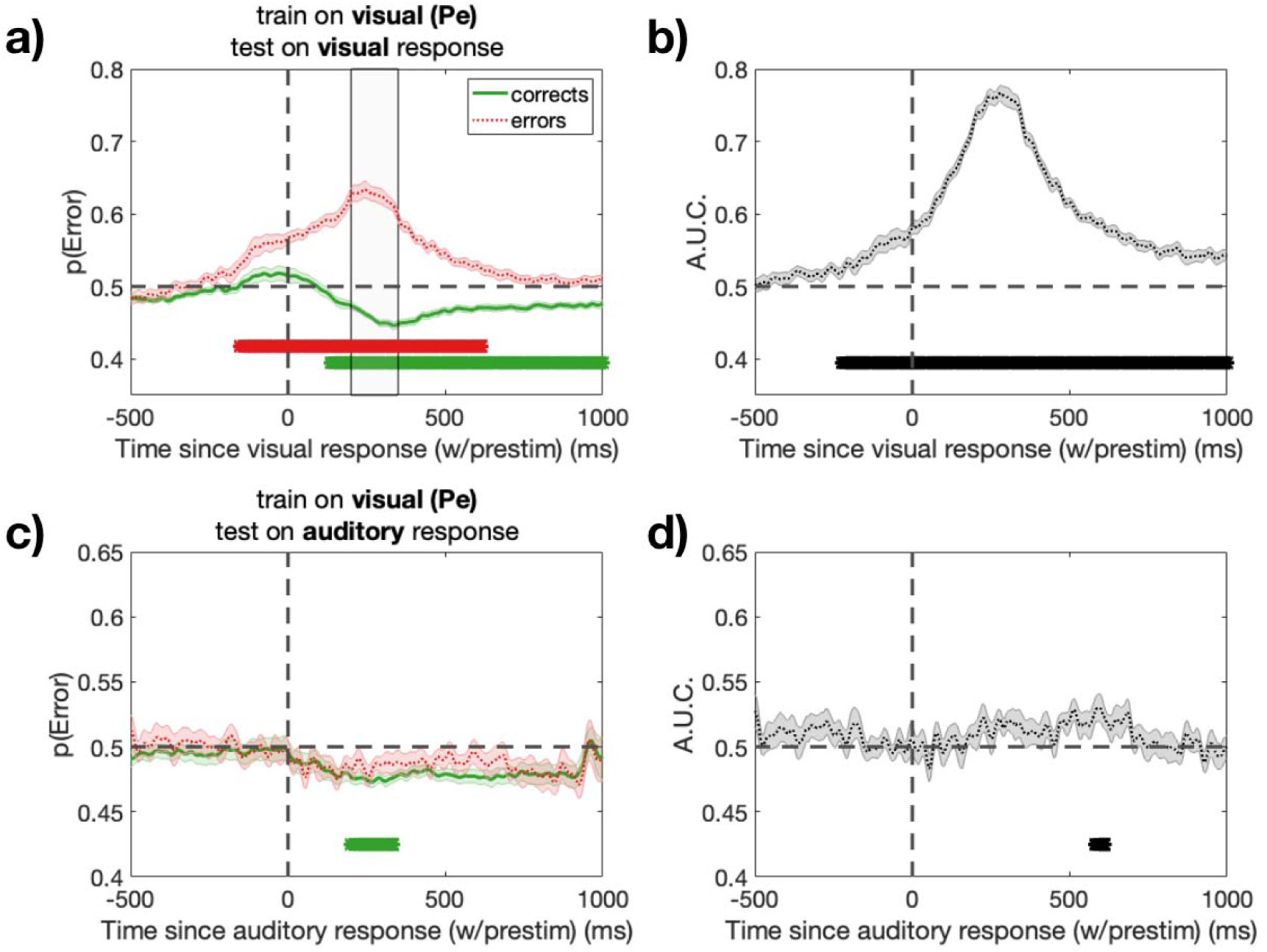
Temporal and crossmodal generalisation of a classifier trained to predict errors on a visual perceptual decision when using response locked data with a pre-stimulus baseline. All conventions as in **Figure 4**.

## References

Bahrami, B., Olsen, K., Latham, P. E., Roepstorff, A., Rees, G., & Frith, C. D. (2010). Optimally interacting minds. Science, 329(5995), 1081–1085.

Baird, B., Smallwood, J., Gorgolewski, K. J., & Margulies, D. S. (2013). Medial and lateral networks in anterior prefrontal cortex support metacognitive ability for memory and perception. The Journal of Neuroscience: The Official Journal of the Society for Neuroscience, 33(42), 16657–16665.

Boldt, A., & Yeung, N. (2015). Shared neural markers of decision confidence and error detection. The Journal of Neuroscience: The Official Journal of the Society for Neuroscience, 35(8), 3478–3484.

Charles, L., & Yeung, N. (2019). Dynamic sources of evidence supporting confidence judgments and error detection. Journal of Experimental Psychology. Human Perception and Performance, 45(1), 39–52.

de Gardelle, V., & Mamassian, P. (2014). Does confidence use a common currency across two visual tasks? Psychological Science, 25(6), 1286–1288.

Desender, K., Boldt, A., & Yeung, N. (2018). Subjective confidence predicts information seeking in decision making. Psychological Science, 29(5), 761–778.

Double, K. S., & Birney, D. P. (2019). Reactivity to Measures of Metacognition. Frontiers in Psychology, 10, 2755.

Falkenstein, M. (1990). Effects of errors in choice reaction tasks on the ERP under focused and divided attention. Psychophysiological Brain Research. https://cir.nii.ac.jp/crid/1574231874973732352

Fernandez-Vargas, J., Tremmel, C., Valeriani, D., Bhattacharyya, S., Cinel, C., Citi, L., & Poli, R. (2021). Subject- and task-independent neural correlates and prediction of decision confidence in perceptual decision making. Journal of Neural Engineering, 18(4). 10.1088/1741-2552/abf2e4

Feuerriegel, D., Murphy, M., Konski, A., Mepani, V., Sun, J., Hester, R., & Bode, S. (2022). Electrophysiological correlates of confidence differ across correct and erroneous perceptual decisions. NeuroImage, 259, 119447.

Fleming, S. M., & Frith, C. D. (2014). *The Cognitive Neuroscience of Metacognition* (S. M. Fleming & C. D. Frith (eds.)). Springer Berlin Heidelberg.

Fleming, S. M., Ryu, J., Golfinos, J. G., & Blackmon, K. E. (2014). Domain-specific impairment in metacognitive accuracy following anterior prefrontal lesions. Brain: A Journal of Neurology, 137(Pt 10), 2811–2822.

Gherman, S., & Philiastides, M. G. (2015). Neural representations of confidence emerge from the process of decision formation during perceptual choices. NeuroImage, 106, 134–143.

Grützmann, R., Endrass, T., Klawohn, J., & Kathmann, N. (2014). Response accuracy rating modulates ERN and Pe amplitudes. Biological Psychology, 96, 1–7.

Henmon, V. A. C. (1911). The relation of the time of a judgment to its accuracy. Psychological Review, 18(3), 186–201.

Herding, J., Ludwig, S., von Lautz, A., Spitzer, B., & Blankenburg, F. (2019). Centro-parietal EEG potentials index subjective evidence and confidence during perceptual decision making. NeuroImage, 201, 116011.

Hillyard, S. A., Squires, K. C., Bauer, J. W., & Lindsay, P. H. (1971). Evoked potential correlates of auditory signal detection. Science, 172(3990), 1357–1360.

Jordano, M. L., & Touron, D. R. (2018). How often are thoughts metacognitive? Findings from research on self-regulated learning, think-aloud protocols, and mind-wandering. Psychonomic Bulletin & Review, 25(4), 1269–1286.

Kepecs, A., & Mainen, Z. F. (2012). A computational framework for the study of confidence in humans and animals. Philosophical Transactions of the Royal Society of London. Series B, Biological Sciences, 367(1594), 1322–1337.

Killingsworth, M. A., & Gilbert, D. T. (2010). A wandering mind is an unhappy mind. Science, 330(6006), 932.

King, J.-R., & Dehaene, S. (2014). Characterizing the dynamics of mental representations: the temporal generalization method. Trends in Cognitive Sciences, 18(4), 203–210.

Kornell, N., Son, L. K., & Terrace, H. S. (2007). Transfer of metacognitive skills and hint seeking in monkeys. Psychological Science, 18(1), 64–71.

Lee, D. G., Daunizeau, J., & Pezzulo, G. (2023). Evidence or Confidence: What Is Really Monitored during a Decision? Psychonomic Bulletin & Review, 30(4), 1360–1379.

Maris, E., & Oostenveld, R. (2007). Nonparametric statistical testing of EEG- and MEG-data. Journal of Neuroscience Methods, 164(1), 177–190.

Masset, P., Ott, T., Lak, A., Hirokawa, J., & Kepecs, A. (2020). Behavior- and Modality-General Representation of Confidence in Orbitofrontal Cortex. Cell, 182(1), 112– 126.e18.

Mazancieux, A., Fleming, S. M., Souchay, C., & Moulin, C. J. A. (2020). Is there a G factor for metacognition? Correlations in retrospective metacognitive sensitivity across tasks. Journal of Experimental Psychology. General, 149(9), 1788–1799.

Morales, J., Lau, H., & Fleming, S. M. (2018). Domain-General and Domain-Specific Patterns of Activity Supporting Metacognition in Human Prefrontal Cortex. The Journal of Neuroscience: The Official Journal of the Society for Neuroscience, 38(14), 3534– 3546.

Pescetelli, N., Hauperich, A.-K., & Yeung, N. (2021). Confidence, advice seeking and changes of mind in decision making. Cognition, 215, 104810.

Petrusic, W. M., & Baranski, J. V. (2003). Judging confidence influences decision processing in comparative judgments. Psychonomic Bulletin & Review, 10(1), 177–183.

Pleskac, T. J., & Busemeyer, J. R. (2010). Two-stage dynamic signal detection: a theory of choice, decision time, and confidence. Psychological Review, 117(3), 864–901.

Qiu, L., Su, J., Ni, Y., Bai, Y., Zhang, X., Li, X., & Wan, X. (2018). The neural system of metacognition accompanying decision-making in the prefrontal cortex. PLoS Biology, 16(4), e2004037.

Rabbitt, P. M. (1968). Three kinds of error-signalling responses in a serial choice task. The Quarterly Journal of Experimental Psychology, 20(2), 179–188.

Rausch, M., Zehetleitner, M., Steinhauser, M., & Maier, M. E. (2020). Cognitive modelling reveals distinct electrophysiological markers of decision confidence and error monitoring. NeuroImage, 218, 116963.

Rouault, M., Lebreton, M., & Pessiglione, M. (2023). A shared brain system forming confidence judgment across cognitive domains. Cerebral Cortex, 33(4), 1426–1439.

Scheffers, M. K., & Coles, M. G. H. (2000). Performance monitoring in a confusing world: error-related brain activity, judgments of response accuracy, and types of errors. Journal of Experimental Psychology. Human Perception and Performance, 26(1), 141–151.

Silver, I., Mellers, B. A., & Tetlock, P. E. (2021). Wise teamwork: Collective confidence calibration predicts the effectiveness of group discussion. Journal of Experimental Social Psychology, 96, 104157.

Steinhauser, M., & Yeung, N. (2010). Decision processes in human performance monitoring. The Journal of Neuroscience: The Official Journal of the Society for Neuroscience, 30(46), 15643–15653.

Su, J., Jia, W., & Wan, X. (2022). Task-Specific Neural Representations of Generalizable Metacognitive Control Signals in the Human Dorsal Anterior Cingulate Cortex. The Journal of Neuroscience: The Official Journal of the Society for Neuroscience, 42(7), 1275–1291.

Valeriani, D., Cinel, C., & Poli, R. (2017). Group Augmentation in Realistic Visual-Search Decisions via a Hybrid Brain-Computer Interface. Scientific Reports, 7(1), 7772.

van den Berg, R., Zylberberg, A., Kiani, R., Shadlen, M. N., & Wolpert, D. M. (2016). Confidence Is the Bridge between Multi-stage Decisions. Current Biology: CB, 26(23), 3157–3168.

Vickers, D., & Packer, J. (1982). Effects of alternating set for speed or accuracy on response time, accuracy and confidence in a unidimensional discrimination task. Acta Psychologica, 50(2), 179–197.

Yeung, N., & Summerfield, C. (2012). Metacognition in human decision-making: confidence and error monitoring. Philosophical Transactions of the Royal Society of London. Series B, Biological Sciences, 367(1594), 1310–1321.

Yu, S., Pleskac, T. J., & Zeigenfuse, M. D. (2015). Dynamics of postdecisional processing of confidence. Journal of Experimental Psychology. General, 144(2), 489–510.

